# Crossmodal Expectations in Material Perception

**DOI:** 10.64898/2026.06.24.734160

**Authors:** Amna Malik, Linda Kölmel, Jutta Billino, Katja Doerschner

**Author notes:** Correspondence to: Amna Malik Postdoctoral researcher, Department of Psychology, Justus Liebig University Giessen, Otto-Behaghel Straße 10F, 35394, Giessen, Germany.

## Abstract

Humans rely on multiple sensory modalities, such as vision, audition, and touch, to perceive materials in everyday life. Previous research shows that multisensory perception leads to facilitation, yet the mechanisms responsible for this facilitation remain poorly understood. One potential mechanism is crossmodal prediction, whereby input from one modality generates predictions about another. While substantial research on multisensory facilitation has focused on bottom-up processes, such as spatial, temporal, and semantic congruency, the role of crossmodal predictions, particularly in material perception, has received little attention. To address this gap, we conducted two experiments, a reaction time task and a material rating task, in which participants viewed computer-generated animations of familiar objects being dropped to the ground. The paradigm exploited the natural temporal structure of impact events: pre-impact visual appearance provides information about an object’s material and therefore can generate expectations about the forthcoming impact sound. Critically, participants saw the event only until before the impact, after which the video was masked. Thus, vision and audition were temporally aligned but not presented concurrently, allowing us to isolate the influence of visually driven expectations on the incoming auditory information without a bottom-up conflict. In some trials, the sound matched the expected material, but in a subset, it was incongruent, violating expectations elicited by the preceding visual information. Across both experiments, participants took longer to respond on incongruent than congruent trials, suggesting increased processing demands. In the rating task, incongruent trials also shifted material judgments, such that ratings reflected a weighted combination of incoming auditory information and visually driven predictions, with large individual differences in relative cue weighting. These findings suggest that priors on material properties from one modality, specifically vision, not only establish high-level expectations within the modality about an object’s future state, but also extend across modalities.

## Introduction

Imagine you’re sitting in a busy cafe, when suddenly you see a waiter lose grip on a large metal serving tray stacked with ceramic plates. Before any sound reaches you, you reflexively tense your shoulders, raising your hands to shield your ears because you anticipate the piercing crash of shattering ceramic. What enables such a preemptive response?

Over repeated interactions with everyday objects, we learn to associate objects with the properties of the materials that they are typically made of (Fleming, 2017; Fleming et al., 2013). These learned associations enable us to form expectations about the future state of an object under external forces, guiding our interaction with it (Alley et al., 2020; Malik et al., 2023). A particularly clear demonstration of such expectation effects comes from the ‘surprising materials’ paradigm introduced by Alley et al. (2020), in which objects were shown falling towards the ground, and observers could strongly anticipate the likely material outcome from the object’s identity and appearance (The complete stimulus set is available on Zenodo: https://doi.org/10.5281/zenodo.2542577). They created expectation violations through ‘surprising’ object-material combinations, in which a familiar object exhibited an atypical material behavior (e.g., a teacup wrinkling as if it were fabric). Under these conditions, perceptual judgments were systematically biased toward the expected outcome, highlighting a strong influence of prior knowledge on material perception. Such expectation violations also led to measurable behavioral costs, including longer response times, consistent with the need to resolve a mismatch between prediction and sensory evidence (Alley et al., 2020). Building on this, Malik et al. (2023) demonstrated a similar cost in a relatively low-level perceptual decision task, showing that expectation violations delayed responses not only for long-term, real-world material associations, but also for newly learned object-material associations acquired within the experiment.

Expectation violation paradigms have been used extensively in perceptual research (de Lange et al., 2018; Summerfield & Egner, 2009). The material expectation paradigms described above, however, differ somewhat from more canonical paradigms in which cues establish expectations about an event’s category, location, or timing. Specifically, they recruit long-term priors grounded in real-world knowledge about the material properties of objects, thereby enabling predictions about how an object will behave under force in a relatively naturalistic setting (Alley et al., 2020). Importantly, the studies discussed above focused on vision, whereas real-world perception is inherently multisensory. (Ernst & Bülthoff, 2004; Shams & Seitz, 2008). Events usually generate signals across multiple modalities that are spatially and temporally aligned and carry meaningful semantic relationships (Ernst & Bülthoff, 2004; Soto-Faraco et al., 2019; Spence, 2011). Crucially, the very kinematic interactions that reveal how an object behaves under force are often accompanied by characteristic sounds. Glass produces a high-pitched shatter when it breaks, wood snaps with a dull crack, and water splashes when perturbed. These auditory signatures are highly diagnostic of the underlying material (Fujisaki et al., 2014; Giordano & McAdams, 2006; Klatzky et al., 2000). Therefore, through repeated exposure, we learn not only how materials deform and move on impact, but also learn these crossmodal audiovisual associations that link the visual appearance of a material with its characteristic sounds. Such associations could enable us to generate predictions across modalities; for example, in the cafe scenario above, you not only anticipate the event’s visual consequences but also its auditory outcome: a high-pitched shattering sound. Here, we ask whether expectations, generated from crossmodal associations, modulate material perception.

A large body of work on multisensory perception has investigated how spatial and temporal congruency shapes audiovisual integration, but they often use simple laboratory stimuli, brief flashes, and tone bursts that lack the kind of semantic associations present in real-world events (Alais & Burr, 2004; Dixon & Spitz, 1980; Meredith et al., 1987; Senkowski et al., 2007; Shams et al., 2002; Vroomen et al., 2004). As a result, the contribution of semantic congruency to multisensory interactions has been comparatively underexplored. Research suggests that semantically congruent audiovisual stimuli, compared to incongruent, can yield faster reaction times in detection and classification tasks (Laurienti et al., 2004; Roberts et al., 2024; Yu et al., 2022; Zhou et al., 2023) and improve target detection in visual search (Iordanescu et al., 2008; Mishra & Gazzaley, 2012). However, in these paradigms, the auditory and visual signals are presented simultaneously. Under these conditions, performance benefits can often be explained, at least in part, by redundant-signals mechanisms such as statistical facilitation (race models), multisensory coactivation, and automatic crossmodal attentional spread (Colonius & Diederich, 2006; Crosse et al., 2015; Miller, 1982; Steinweg & Mast, 2017). In naturalistic settings, observers not only integrate concurrent inputs from multiple modalities but also continuously generate crossmodal predictions about forthcoming events. Thus, while classic congruency paradigms have been invaluable for characterizing online multisensory interactions, they provide limited insight into how top-down crossmodal predictive mechanisms shape multisensory perception. Some sound-to-symbol matching paradigms do probe crossmodal prediction effects. Participants learn arbitrary mappings between simple auditory tokens (tones/beeps) and visual symbols (icons/shapes), and a sound signals which visual target is likely to appear (Dercksen et al., 2021; Pieszek et al., 2013; Stuckenberg et al., 2019; Widmann et al., 2004). These studies demonstrate predictive benefits across modalities, but they remain constrained to highly simplified, semantically sparse stimuli and artificial associations. By contrast, work employing complex, semantically rich stimuli has focused mostly on speech (Peelle & Sommers, 2015; Sohoglu et al., 2012), where predictive mechanisms linking acoustic cues, lexical context, and meaning are well established. Outside of speech, predictive crossmodal interactions with naturalistic events are rarely studied.

To address this research gap, we used video animations of falling objects from Alley et al. (2020) and created a crossmodal expectation violation stimulus set by pairing each video with either a matched (congruent) or mismatched (incongruent) impact sound, extending their logic of “surprising” object-material combinations to the audiovisual domain. Crucially, the visual and auditory streams were temporally aligned but not presented simultaneously: the visual sequence was masked at impact while the sound occurred at that same moment. This eliminated classic bottom-up audiovisual conflict and instead enabled a direct test of whether expectations about material properties derived from preceding visual information influence the interpretation of forthcoming auditory information. The design also mirrored everyday perception, where visual input often precedes sound and provides a temporal window for prediction. We then conducted two experiments: a reaction-time 2-AFC task and a material-rating task. The reaction time task, following Malik et al. (2023), provided a relatively low-level behavioral measure of expectation violation by using a simple yes/no decision to test whether incongruent conditions produce an immediate processing cost, expressed as slower responses. The material rating task assessed how expectation violations shape perceptual judgments by measuring perceived material qualities across multiple attributes and comparing ratings between congruent and incongruent audiovisual pairs. Across both experiments, participants took longer to respond on incongruent than on congruent trials, suggesting greater processing demands to resolve the conflict between visual prior and incoming crossmodal sensory evidence when expectations were violated. In the rating task, incongruent trials showed shifted material judgments, such that ratings reflected a weighted combination of incoming auditory information and visually driven predictions, with some individuals relying more on vision and others more on audition. These findings suggest that priors from one modality, specifically vision, not only establish high-level expectations within the modality about an object’s future state, but also extend across modalities. These crossmodal expectations may serve as a possible mechanism for multisensory facilitation in material perception.

## Materials and Methods

### Experiment 1- 2-AFC reaction time task

In Experiment 1, we used a simple 2-AFC reaction-time task to quantify congruency effects with minimal response demands, avoiding additional delays associated with continuous rating responses (e.g., moving a slider).

#### Participants

Forty-five participants (14 male and 31 female, mean age: 24.8 years, range: 19-82 years) participated in the experiment. All participants had no uncorrected visual or hearing impairments (self-report). Participants provided written informed consent prior to the experiment. Experimental procedures were approved by the ethics board at Justus-Liebig University Giessen and were carried out in accordance with the guidelines outlined in the Declaration of Helsinki. No direct remuneration was paid; instead, ten vouchers were awarded by random draw.

#### Stimuli

We used video animations from Alley et al. (2020) depicting objects dropped from a fixed height and interacting with the ground in ways characteristic of their material properties. From the full stimulus set, we selected only animations of familiar objects (15 in total). These stimuli comprised five material categories: shattering, splashing, nondeforming, wobbling, and wrinkling, with three animations per category summarized in Supplementary Table 1. Each animation consisted of 48 frames, and the object contacted the ground on the 12th frame. To prevent observers from viewing the post-impact event, all frames from the moment of impact (frame 12) to the end of the animation were replaced with a uniform grey mask. The mask luminance corresponded to the mean luminance averaged across all frames of all animations.

Although observers never saw the post-impact outcome, they heard an impact sound matched to the acoustic signature of each material. These sounds were produced by a contracted sound designer for our specific stimulus set, using materials selected to yield clear and unambiguous impact sounds (see Supplementary Table 1). Recordings were made using two Lavalier microphones (DPA 4060) mounted on a modified microphone stand, in combination with an NTG 4+ shotgun microphone (RODE Microphones) and a MixPre-6 recorder (Sound Devices, LLC). The recording setup is shown in Supplementary Figure S1. To generate the sounds, the objects and materials were dropped into a custom-built basin made of coated MDF, and the resulting audio was recorded. The recordings were then edited and temporally aligned to the “frame of impact” in the videos (frame 12). Audio conversion and editing were performed in Reaper (Cockos Inc.), and recordings were processed using a Pro-Q 3 equalizer (FabFilter) and Spectral DeNoise (iZotope, Inc.) for noise reduction.

To reproduce the “surprising” object-material combinations from Alley et al. (2020), we applied the same mapping between visual object identity and unexpected material behavior reported in their study and paired each video with an incongruent impact sound corresponding to the unexpected material class, resulting in 15 incongruent audiovisual stimuli (Supplementary Table 2). Audio tracks (.wav) were combined with the videos in Blender (v3.3.0) to generate .mp4 movies (Blender Online Community, 2024). An overview of the animation sequence with representative frames is shown in Figure 1.

**Figure 1.**
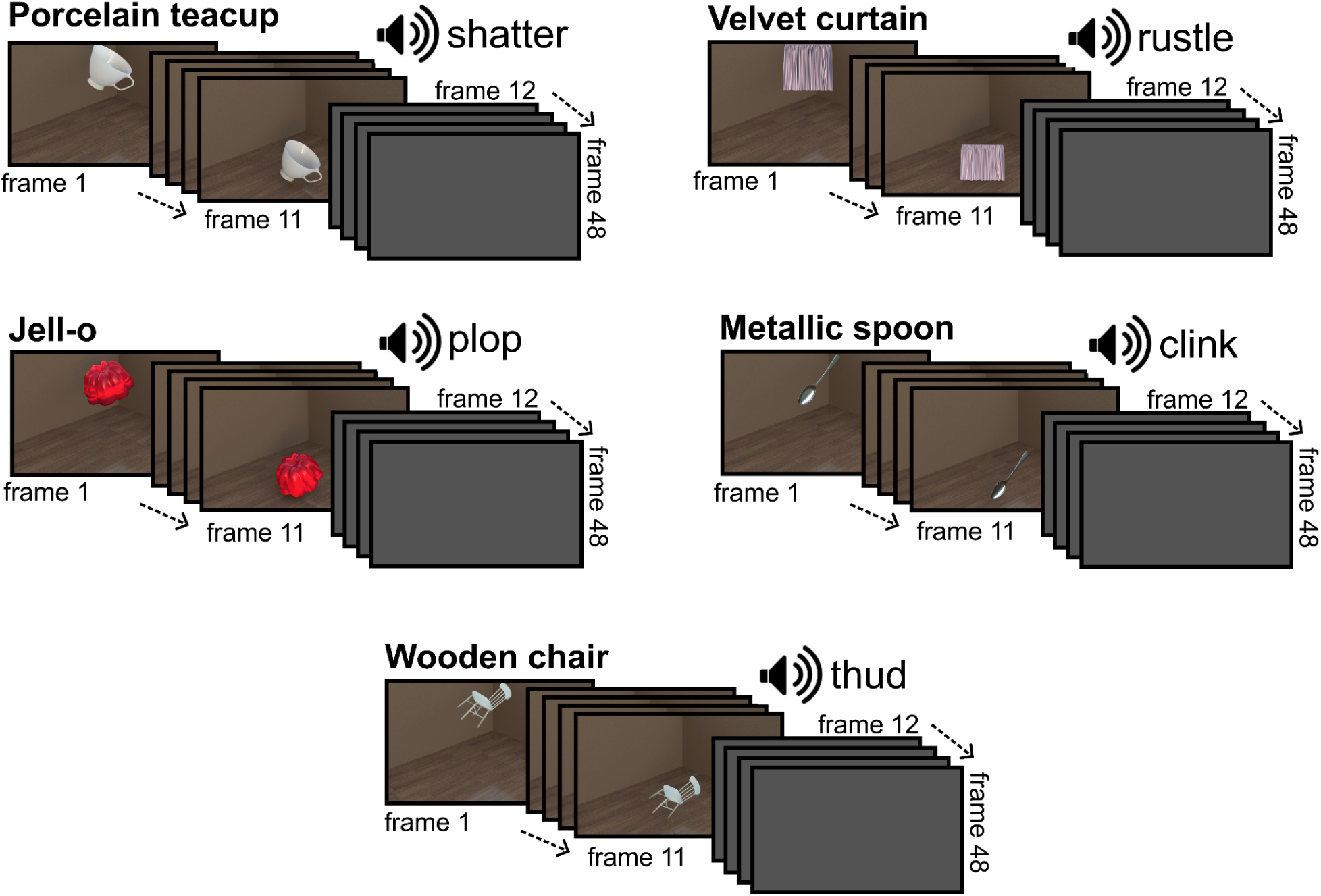
Set of congruent animations used in the current study. The object contacted the ground on frame 12; from this point onward (frames 12-48), the video was replaced with a uniform grey mask, so the impact itself and the post-impact outcome were visually hidden. The speaker icon indicates the characteristic (congruent) impact sound for each animation, which began at the moment of impact (frame 12).

#### Experimental setup

The experiment was conducted on a mobile setup on a laptop computer (Lenovo ThinkPad; 16-inch display, Windows 10), allowing data collection to take place in different testing locations that were convenient and easily accessible for participants. Stimuli were presented at a 60 Hz refresh rate with a screen resolution of 1920 × 1080 pixels. Testing was conducted in a quiet environment with the laptop placed on a table and participants seated on a chair. The viewing distance was approximately arm’s length. Sounds were presented via noise-cancelling headphones (Bose QuietComfort 35 II) with the volume set to 40%, which also helped minimize distracting background noise during the experiment. Participants provided their responses using two handheld custom made response grips. To ensure that participants consistently remembered the response-hand mapping, two laminated A4 signs labeled ‘Yes’ and ‘No’ were placed next to the laptop. Stimuli were presented using the Psychtoolbox (Brainard, 1997) on MATLAB version 2023b (MathWorks Inc., Natick, MA, USA).

#### Procedure

Participants were shown the setup and received standardized instructions. No information about the purpose of the experiment was given in order to avoid biasing perception, and instructions were phrased neutrally (e.g., “an object will be shown” and “a sound will be presented”). Participants were instructed to respond only after the object contacted the ground. If asked whether they needed to respond quickly, they were told that speed was not emphasized and that they should answer the question (“Broken or not?”) as intuitively as possible.

A schematic of a single trial is shown in Figure 2A. Each trial began with the presentation of a fixation cross (1s), followed by the first frame of the animation presented for 1s, providing a static preview of the object. The animation then began playing at the rate of 24 frames per second for 2.0 s, with audio sampled at 48 kHz. At the 11th frame (≈458 ms), which corresponded to the moment of impact, the video was replaced with a uniform gray mask. Importantly, because the point of impact was sharply defined in every animation, the accompanying audio always began precisely when the video was masked, with its duration depending on the physical event being depicted. Participants then indicated whether the object broke upon impact (“Yes”) or did not break (“No”). Each response immediately initiated the next trial. The experiment lasted approximately 2-3 minutes in total.

**Figure 2.**
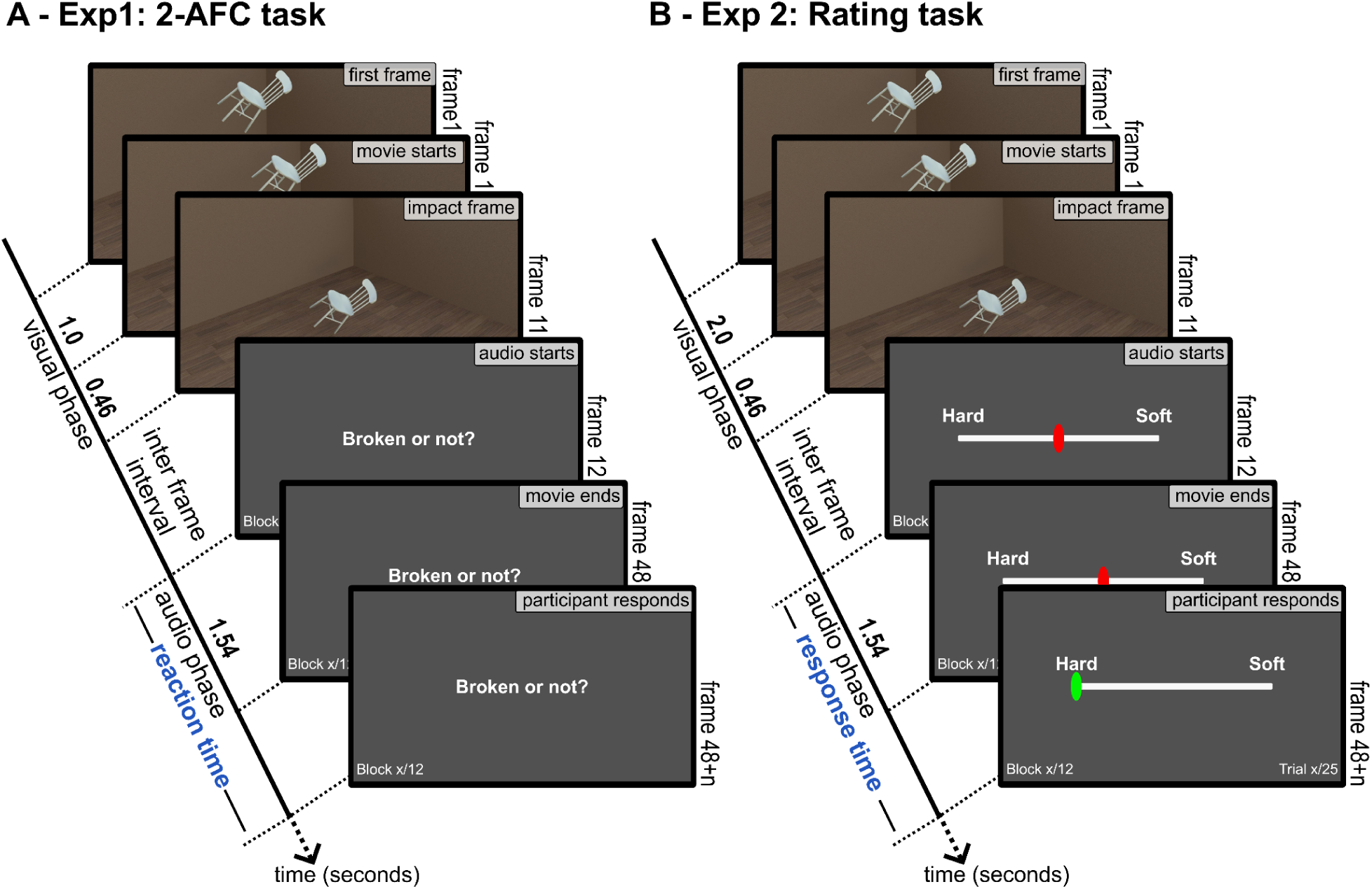
Schematic of a single trial for A) Experiment 1: 2-AFC reaction time task, and B) Experiment 2: Rating task. Note that there was a fixation screen for 1 s before the visual phase.

#### Analysis

Reaction times (RTs) were measured from sound onset (i.e., the moment of impact) to the keypress. Trials with non-positive RTs or RT < 300ms (*n* = 56) or RTs outside ± 2 SD from the mean (*n* = 47) were excluded leaving 1277 trials. Because raw RTs were positively skewed, we applied a Box-Cox transformation, which improved normality and reduced skew (as assessed by visual inspection of histograms and Q-Q plots). We then, fitted a linear mixed-effects model using the lme4 package (Bates et al., 2015) with reaction time as the dependent variable, congruency (congruent vs. incongruent) as a fixed effect, and a random intercept for each participant:

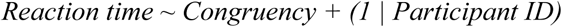

### Experiment 2- Material rating task

We ran Experiment 2 to test the perceptual consequences of crossmodal expectation violations: specifically, whether an incongruent impact sound also affects the perceived material qualities, as reported in the corresponding unimodal design in Alley et al. (2020). Using attribute ratings, we quantify how participants integrate visually driven predictions and incoming auditory cues.

#### Participants

A separate group of 35 participants (12 male and 23 female, mean age: 26.17, range: 20–38 years) took part in Experiment 2. All participants had normal or corrected-to-normal vision and hearing (self-report). Participants provided written informed consent prior to the experiments. Experimental procedures were approved by the ethics board at Justus-Liebig University Giessen and were carried out in accordance with the guidelines outlined in the Declaration of Helsinki. Participants were compensated at the rate of 8 euros per hour.

#### Stimuli

Stimuli specifications were exactly the same as Experiment 1. But for Experiment 2, we selected a subset of five familiar objects, each representing a distinct material category: porcelain teacup, metal spoon (nondeforming & metal), velvet curtain (wrinkling), Jell-O mold (wobbling), and wooden chair (nondeforming & wood), as shown in Figure 1. Each video was paired with its corresponding impact sound to form five congruent audiovisual stimuli (shatter, clink, rustle, plop, and thud, respectively). To create the incongruent stimulus set, each of the five videos was paired with the four remaining sounds, yielding 20 incongruent combinations (5 × 4). Audio tracks (.wav) were combined with the videos in Blender v3.3.0 to generate .mp4 movies (Blender Online Community, 2024).

The stimulus set was reduced to keep the material-rating task manageable and non-exhaustive, while still allowing us to test a fully crossed incongruency design (i.e., all non-matching sound pairings for each visual event), rather than restricting incongruency to a small subset of “surprising” pairings as in Alley et al. (2020). The five selected animations were chosen because they exhibit clearly distinguishable material behaviors and highly diagnostic impact sounds, providing strong separation along the material attributes assessed in the rating task (see below).

#### Experimental setup

Stimuli were presented on a 27-inch LCD color monitor (Dell UltraSharp U2723QE) with a refresh rate of 60 Hz and a resolution of 3840 × 2160 pixels, controlled by a Dell system running on Windows 11. The experiment was conducted in a dark room, with the monitor serving as the only light source. A chinrest was used to minimize head movement at a distance of 60 cm from the screen. Sounds were presented via noise-cancelling headphones (Sennheiser ACCENTUM) with the volume set to 70 %. Stimuli were presented using Psychtoolbox (Brainard, 1997) on MATLAB version 2024b (MathWorks Inc., Natick, MA, USA).

#### Procedure

Before the main experiment, participants received instructions and completed a short practice session with a separate set of stimuli consisting only of congruent pairs. This familiarized them with the task without being exposed to the experimental stimuli.

Each trial began with a 2-s static preview of the object (first frame). The animation then started, and stimulus presentation and timing were otherwise comparable to Experiment 1: the animation played at 24 fps, and at the moment of impact, the display was replaced by a uniform grey mask. After the impact, a rating scale appeared in the center of the screen. This was presented as a horizontal slider anchored by opposing descriptors (e.g., soft-hard). Participants adjusted a continuous slider to indicate which level best matched their perception. After confirming their selection, the slider turned green and they pressed the space bar to proceed to the next trial. A schematic of a single trial is shown in Figure 2B.

**Rating attributes:** Participants rated each stimulus on four bipolar attributes: breaking/deforming-stays intact, hard-soft, wet-dry, and metallic-non-metallic. All four attributes were inherently multimodal in the sense that both auditory and visual information provided cues to rate the given attribute. For example, hardness could be distinguished not only from how an object deformed visually but also from the characteristic sound it produced upon impact; similarly, wetness, metallicity, and intactness were each supported by diagnostic signals in both modalities. This design ensured that neither audition nor vision alone was sufficient, making crossmodal prediction meaningful. Moreover, the set of objects was selected such that each material occupied a distinct location within this four-dimensional attribute space. No two materials shared the same profile across the four ratings, providing clear separability of material categories.

The experiment consisted of 12 blocks, each focused on a single attribute. Each block contained 25 trials (five congruent, twenty incongruent), and all four attributes were repeated three times across the session. Blocks were arranged in a pseudo-randomized order so that each set of four blocks covered all four attributes before repeating. Two measures were collected on each trial: the position of the slider (attribute rating) and the response time. Response time was defined as the interval between the appearance of the slider (i.e., the impact frame) and the confirmation of the response. On average, the experiment lasted 30 minutes.

#### Analysis

**Response time data:** Response times were measured from sound onset (i.e., the moment of impact) to the keypress. Trials with non-positive RTs, RT<300ms or RTs outside ± 2 SD from the mean were excluded (n = 421), leaving 10,079 trials. Since the distribution of raw response times was not normally distributed, we transformed them using the Box-Cox transformation. To test the effect of congruency, we fitted a linear mixed-effects model using the lme4 package (Bates et al., 2015) with response time as the dependent variable, congruency (congruent vs. incongruent) as a fixed effect, and a random intercept for each participant:

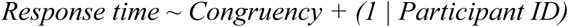

#### Rating Data

**Intra- and inter-observer consistency:** Both intra- and inter-observer reliability were quantified using two-way random-effects ICC(2,1). Intra-observer ICCs captured the stability of each participant’s ratings across repeated trials for congruent and incongruent trials. Inter-observer ICCs assessed agreement across participants by comparing their mean ratings for each condition

#### Comparison of congruent and incongruent ratings

To quantify the effect of preceding visual information on incoming auditory information in incongruent trials, we used congruent audiovisual trials as ecologically valid reference values, because they reflect participants’ judgments when auditory and visual information are naturally consistent. For each incongruent audio-video combination, we defined two reference points: the mean congruent rating for the auditory stimulus in the AV pair (A_ref_) and the mean congruent rating for the visual stimulus (V_ref_) in the AV pair. For example, if the visual stimulus was a teacup and the auditory stimulus was jelly, V_ref_ was the mean rating in the congruent teacup condition (teacup video + teacup sound), whereas A_ref_ was the mean rating in the congruent jelly condition (jelly video + jelly sound).

When visual and auditory cues in an incongruent AV pair suggest similar material qualities (e.g., a visually soft fabric paired with a similarly “soft” sound), ratings are less diagnostic because both cues point in the same direction. In contrast, contrasting pairs (e.g., visually soft + acoustically hard, or vice versa) provide a clearer test of crossmodal bias. Therefore, to make incongruency effects interpretable, we restricted the analysis to maximally contrasting incongruent pairs, in which the visual event and impact sound implied opposite ends of the rated attribute. We encoded this manipulation as *conflict direction*, indicating whether the visual cue corresponded to the high end of the attribute and the auditory cue to the low end (Vhigh-Alow), or vice versa (Vlow-Ahigh). Poles were defined from congruent trials: objects with mean ratings ≤ 0.2 were assigned to the low pole and those with mean ratings ≥ 0.8 to the high pole; intermediate objects (0.2–0.8) were excluded. This categorization helps us test whether visual bias differs depending on whether vision predicts the high vs. low pole of the attribute. Supplementary Table 3 summarizes the resulting stimulus set and labels.

*Prior Pull:* Prior pull quantifies whether ratings on incongruent trials were biased toward the visual reference (V_ref_) rather than the auditory reference (A_ref_). For each incongruent trial with rating R from the above filtered data, we computed a normalized prior pull index:

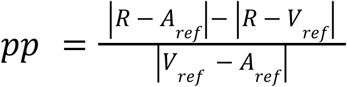

With this definition, PP > 0 indicates that the rating is closer to *V_ref_* (bias toward the visual prior), PP < 0 indicates that it is closer to *A_ref_*(bias toward the auditory cue), and PP = 0 indicates equal proximity to both references. We fitted a linear mixed-effects model with prior pull as the dependent variable, and attribute, conflict direction (Vhigh–Alow vs. Vlow–Ahigh), and their interaction as fixed effects, with random intercepts for participant:

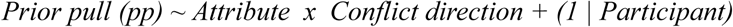

We assessed fixed effects using Type III ANOVA. To follow up significant interactions, we used estimated marginal means for pairwise comparisons, testing (i) differences between conflict directions within each attribute, and (ii) differences in prior pull across attributes (Tukey-adjusted for multiple comparisons where applicable).

## Results

### Experiment 1- Reaction time task

A linear mixed-effects model revealed a significant main effect of Condition (β = 0.112, SE = 0.032, t(1232) = 3.52, p < .001). Reaction times were significantly longer for incongruent trials compared to congruent trials (see Figure 3-A), consistent with the prediction that conflicting audiovisual cues increase processing demands. The effect was small in magnitude (model-based Cohen’s *d* = 0.20).

**Figure 3.**
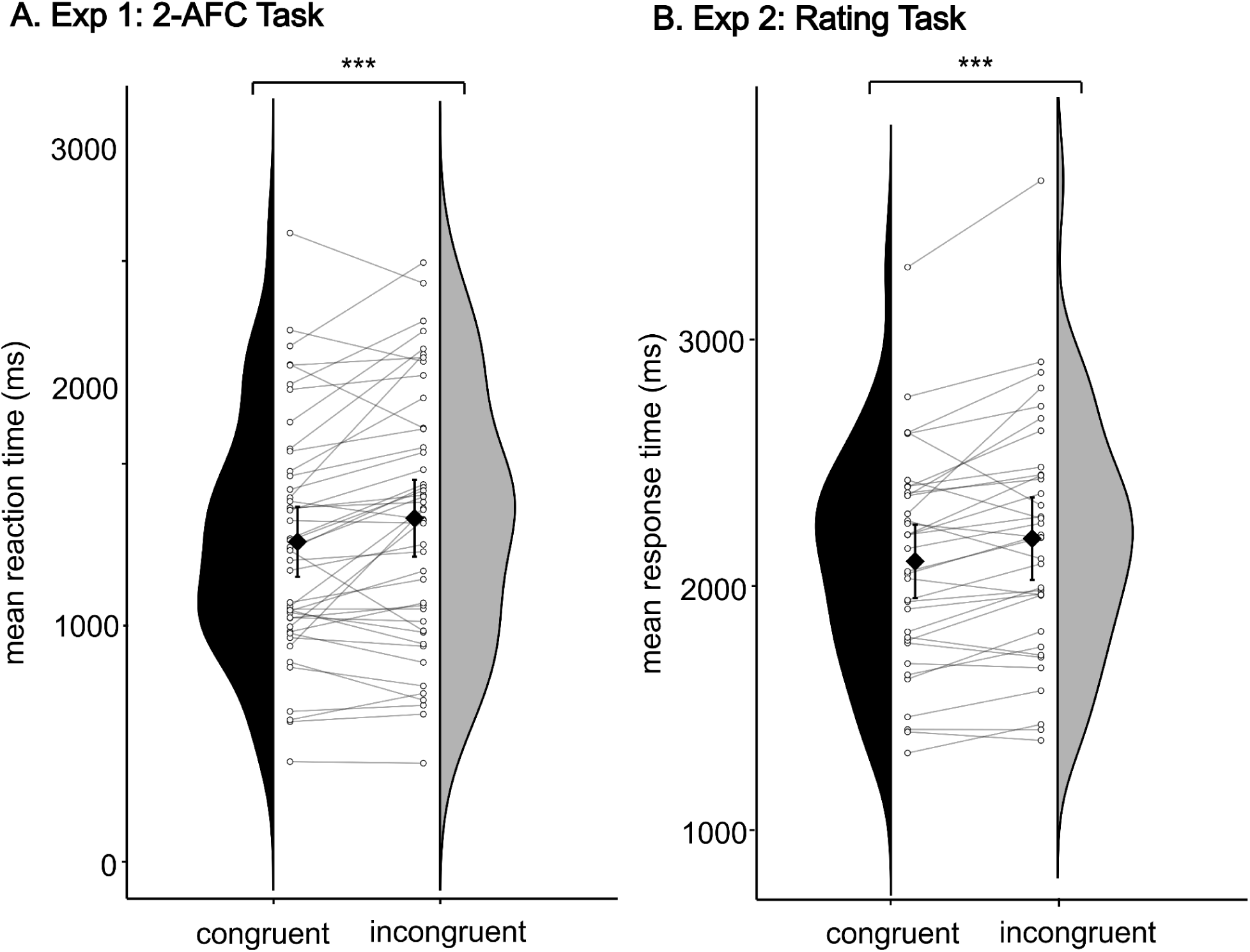
Response times (ms) by audiovisual congruency across two tasks; (A) Two-alternative forced-choice (2AFC) reaction time task. (B) Material rating task. Half-violin plots show the distributions of participants’ mean response times across congruency conditions. White dots indicate individual participants’ mean response times, with lines connecting paired observations across conditions. Black diamonds mark the group means; error bars indicate 95% confidence intervals across participants.Asterisks denote the significance of the congruency contrast (p < .001).

### Experiment 2- Material rating task

**Response time data:** A linear mixed-effects model revealed a significant main effect of Congruency (β = 0.0385, SE = 0.009, t(10,040) = −4.23, p < .001). Similar to Experiment 1, Response times were significantly longer for incongruent audio–video pairs compared to congruent pairs (see Figure 3-B). The effect was small in magnitude (model-based Cohen’s d = −0.11).

#### Rating Data

**Intra- and inter-observer consistency:** Intra-observer consistency was high for both conditions, with ICC values of 0.87 for congruent trials and 0.76 for incongruent trials, indicating that participants were consistent in their own ratings across repetitions. Inter-observer consistency showed a different pattern: in the congruent condition, ICC was 0.75, reflecting strong agreement across participants, whereas in the incongruent condition, ICC dropped to 0.41, indicating considerably weaker agreement (Figure 4). This reduction suggests that, while participants are consistent in their judgements on congruent trials, they show variability on incongruent trials.

**Figure 4.**
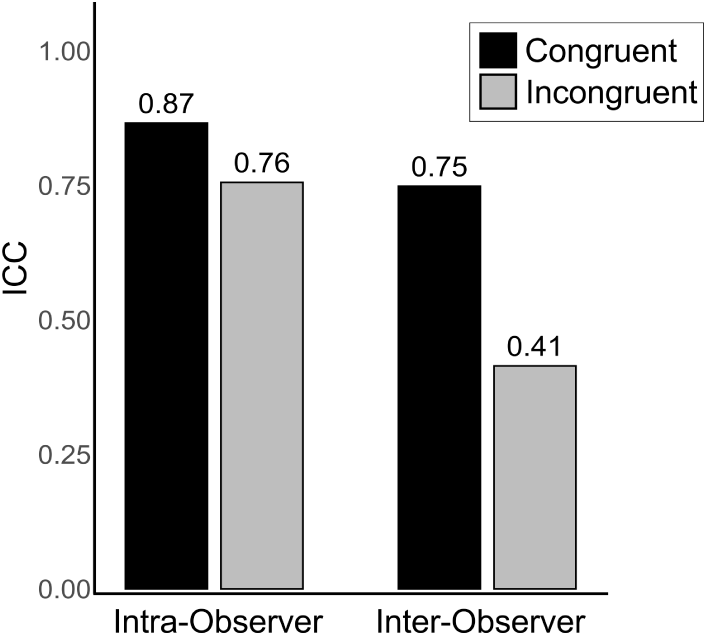
Intra- and inter-observer intraclass correlations (ICCs) for congruent and incongruent conditions.

**Congruent versus incongruent ratings:** Figure 5 shows the mean attribute ratings for congruent animations, together with one representative incongruent condition. In the congruent condition, each material exhibits a distinct and characteristic pattern across attributes, providing a unique perceptual “signature” for that object. The incongruent examples are included as a reference to illustrate how this signature shifts when the impact sound is inconsistent with the expectation generated by the preceding visual information.

**Figure 5:**
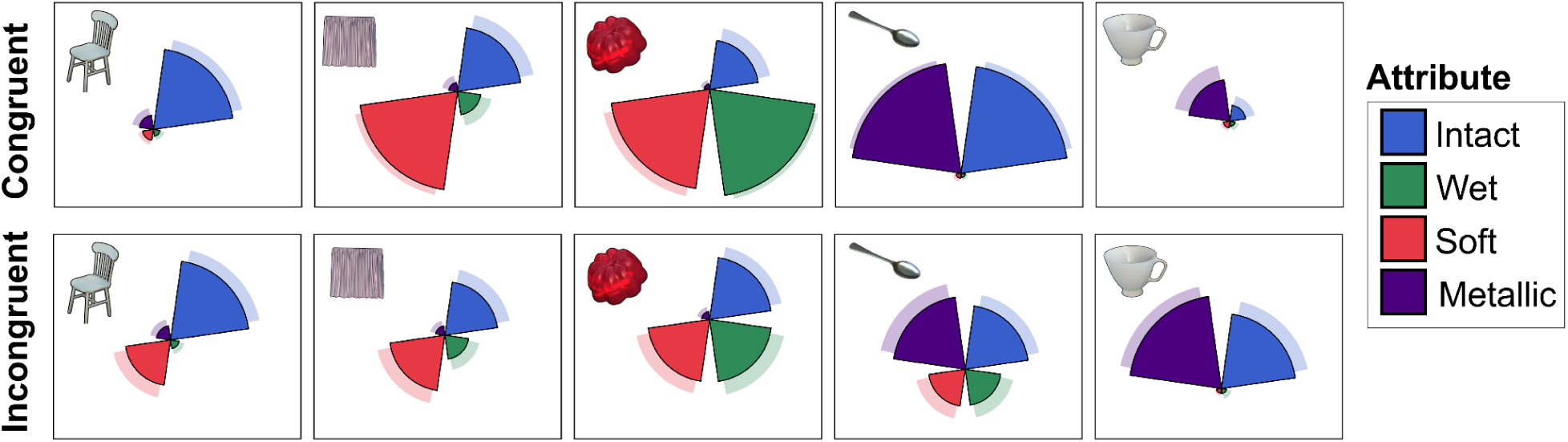
Radar plots of the perceptual profiles for each of the five objects (porcelain teacup, metallic spoon, velvet curtain, Jell-O mold, wooden chair). Top row: congruent audiovisual trials, in which each object is paired with its matching impact sound. Bottom row: example incongruent trials, in which the same impact sound is paired with a visually incongruent object; one representative incongruent pairing is shown for each object.Each axis represents one of four perceptual attributes: softness, wetness, metallicity, and intactness. Greater distance from the center indicates stronger attribution of that property, whereas little or no extension indicates attribution of the opposite pole of that property (i.e., deformability, dryness, hardness, or nonmetallicity). Darker shading indicates the mean rating, and lighter shading indicates the upper 95% confidence bound. Together, these plots illustrate the distinct perceptual “signature” of each material in four dimensional space and how that profile shifts under incongruent condition.

Figure 6 shows the ratings for both congruent and incongruent conditions, comparing each congruent audiovisual pair with all incongruent pairs involving the same sound. In the incongruent condition, when the audio and video cued the same end of an attribute (e.g., a soft video paired with a soft sound), ratings were straightforward. By contrast, when the auditory input cued a conflicting value relative to the preceding video, participants’ ratings reflected a weighted combination of visual priors and auditory information. In other words, the visual prior pulled the average rating toward itself, as illustrated by the red arrows in Figure 6. The raw data underlying these ratings are shown in Supplementary Figures S2 and S3.

**Figure 6:**
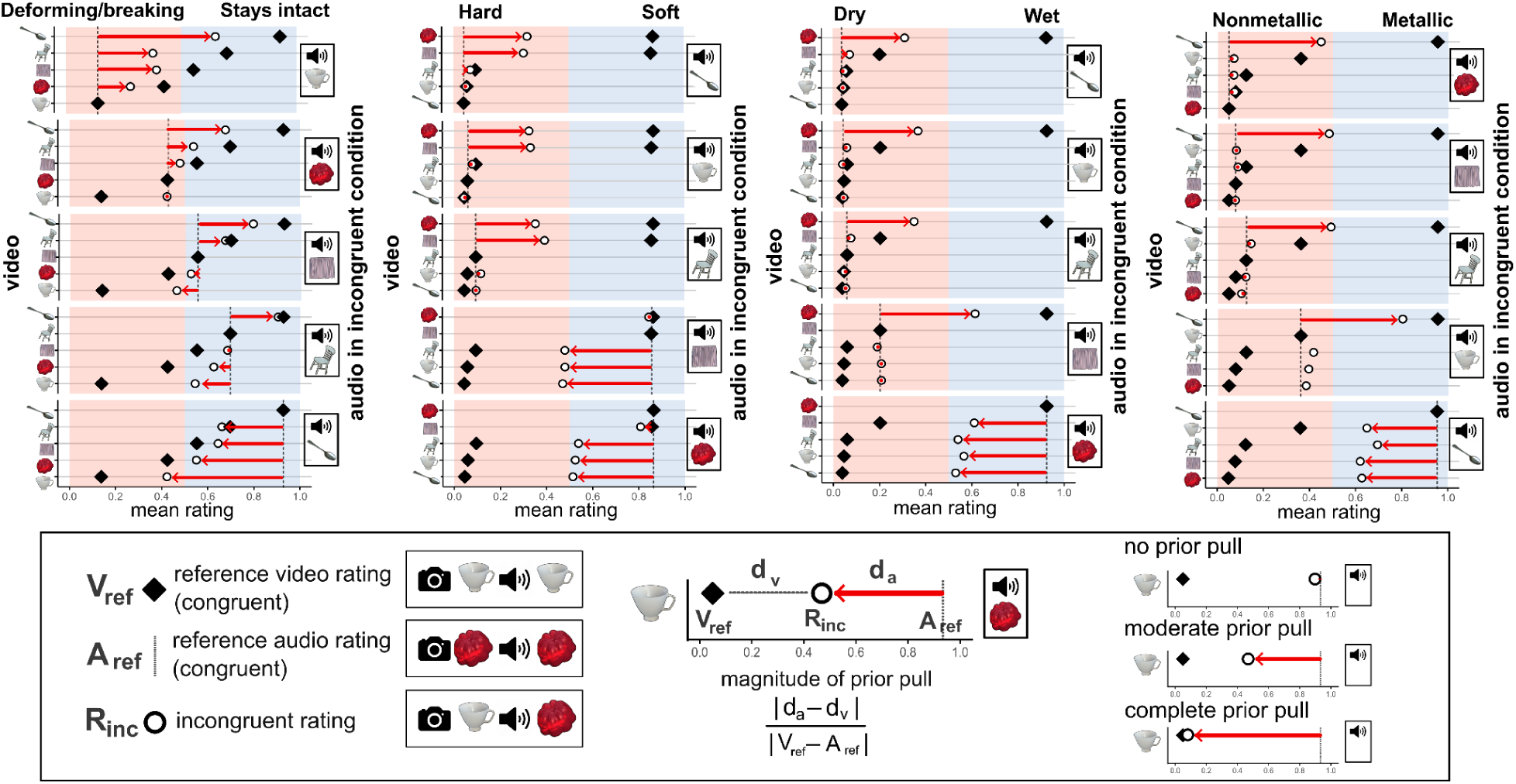
Mean congruent and incongruent ratings for each object across the four perceptual attributes represented by four columns. The x-axis represents the attribute rating scale (0–1). And the y-axis in each plot indicates the reference video from an incongruent audio-video pair, while the black square on the right edge of each plot shows the corresponding reference audio. Black diamonds show the reference video ratings (congruent video–audio pairs), dashed vertical lines show the reference audio ratings (congruent audio–video pairs), and open circles represent the mean incongruent ratings. The blue and red backgrounds depict the contrasting poles of the same attributes. Red arrows illustrate the prior pull, with shifts toward the video reference indicating the influence of visual priors. The schematic (bottom) shows how prior pull was computed: d_v_ is the distance from the incongruent rating to the video reference, d_a_ is the distance from the incongruent rating to the audio reference. The rightmost schematic provides examples of no, moderate, and complete prior pull.

**Prior Pull:** To quantify this shift, we computed a *prior pull index* for contrasting audiovisual incongruent pairs, which measures whether ratings were biased toward the visual reference value or the auditory reference value. This allowed us to test statistically whether, and under which conditions, material judgments were pulled toward the visually induced prior. A linear mixed-effects model with prior pull as the dependent variable revealed a significant main effect of attribute (χ²(3) = 134.91, p < .001) and a significant interaction between attribute and conflict direction (χ²(3) = 66.44, p < .001), but no main effect of Level (χ²(1) = 3.16, p = .075). Estimated marginal means (Figure 7A) showed that prior pull differed reliably between conflict direction for wet-dry, hard-soft, and metallic-nonmetallic (all p < .001), whereas the conflict direction was not different for breaking/deforming-stays intact (p = .162). Across attributes, breaking/deforming-stays intact elicited stronger prior pull than all other attributes (p < .001).

**Figure 7.**
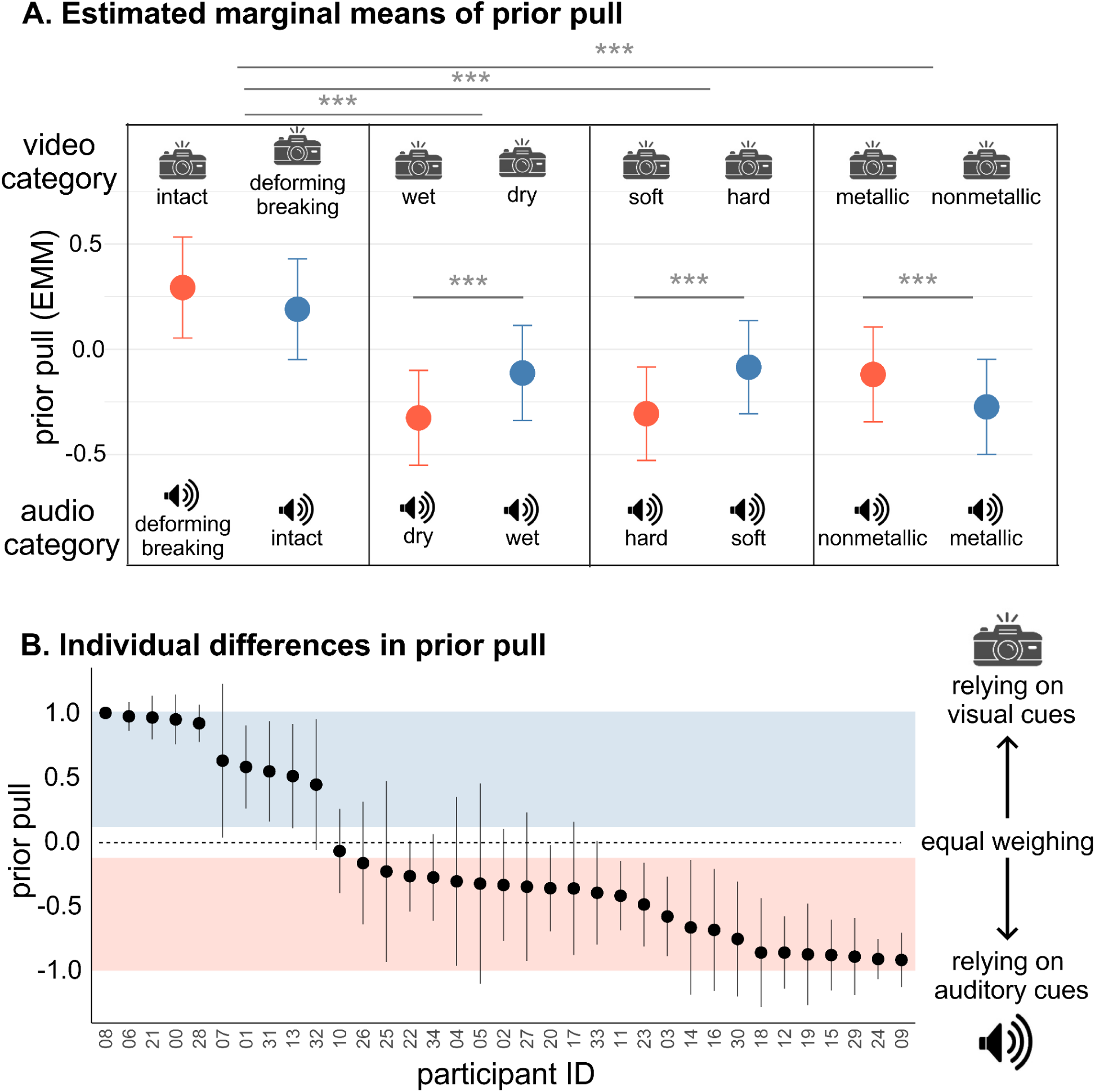
(A) Estimated marginal means of prior pull as a function of attribute and conflict direction. Each point represents model estimates with 95% confidence intervals. (B) Individual differences in prior pull across participants. Each point represents a participant’s mean prior-pull value with error bars indicating ±1 SD. Participants are ordered from highest to lowest mean prior pull. The blue shaded region marks positive prior pull values, indicating greater reliance on preceding visual information when judging material properties. The pink shaded region marks negative prior pull values, indicating greater weighting of incoming auditory information. Values in the intermediate range indicate that judgments reflect the integration of both visual and auditory information.

Participant-to-participant variability in the magnitude of prior pull was substantial, as shown in Figure 7B. Some participants showed predominantly positive prior pull values, indicating greater reliance on prior visual information when judging material properties, whereas others showed predominantly negative prior pull values, indicating greater reliance on incoming auditory information. Many participants’ judgments fell between these extremes, reflecting integration of both visual and auditory information. Consistent with this, the model explained substantial variance overall (conditional R² = 0.60) but little via fixed effects alone (marginal R² = 0.034).

## Discussion

The present study provides evidence that material expectations generated from vision extend across modalities and influence the processing of subsequent auditory input. Critically, by temporally separating visual and auditory signals, our paradigm isolates predictive influences from bottom-up audiovisual integration, suggesting that the observed effects reflect top-down expectations derived from prior visual input rather than concurrent multisensory processing. When these expectations were violated, participants exhibited longer response times, indicating a processing cost associated with resolving conflict between predicted and incoming sensory information.

Beyond response speed, our results further reveal that crossmodal predictions also shape perceptual judgments. Our findings further show that when auditory and visual cues conflict, material judgments are influenced by both sources of information: responses shift toward a weighted combination of the auditory signal and visually derived expectations. Importantly, these weights varied substantially across individuals, pointing to pronounced inter-individual differences in how sensory evidence and prior expectations are integrated across modalities. To our knowledge, this study provides one of the first demonstrations of crossmodal predictive effects in dynamic, naturalistic material events. It extends research on crossmodal prediction beyond simplified laboratory stimuli and beyond highly specific domains such as speech perception, where predictive mechanisms have been more extensively studied. More broadly, these findings highlight the importance of crossmodal predictive processes in shaping multisensory perception under more naturalistic conditions.

### Response times increase when expectations about the material properties of an object are violated across modalities

Across both experiments, participants took longer to respond when the impact sound violated the material expectation elicited by the preceding visual information. In Experiment 1, this effect was expressed in reaction times (RTs) for a speeded yes/no judgment. In Experiment 2, the same pattern was evident in response times for the rating decision. For clarity, we refer to these measures collectively as response times when discussing both experiments together.

While this pattern might at first resemble classic findings that congruent audiovisual stimuli often yield faster responses than incongruent stimuli (Laurienti et al., 2004; Roberts et al., 2024; Yu et al., 2022; Zhou et al., 2023), the underlying explanation is different. Classic congruency effects are frequently discussed in terms of online multisensory facilitation under concurrent stimulation, including mechanisms involving redundant signals (Colonius & Diederich, 2006; Crosse et al., 2015; Miller, 1982; Steinweg & Mast, 2017). Here, congruency was used to probe crossmodal prediction: because visual input was removed at impact while the sound began, the critical mismatch was not between simultaneously presented cues but between a visually elicited expectation and the subsequent auditory evidence in the absence of concurrent audiovisual stimulation. The resulting delays on incongruent trials, therefore, reflect an expectation-violation cost, consistent with prediction-based accounts (De Loof et al., 2016; Urgen & Boyaci, 2021).

A closer conceptual match comes from work that explicitly targets audiovisual prediction, often using symbol-to-sound paradigms in which visual cues predict upcoming sounds (e.g., a symbol predicts a high vs. low tone). Across these paradigms, violated expectations elicit neural mismatch responses such as the mismatch negativity (MMN), an event related potential (ERP) associated with violations of predicted auditory input, and are accompanied by behavioral costs such as slower responses (Dercksen et al., 2021; Pieszek et al., 2013; Stuckenberg et al., 2019; Widmann et al., 2004). While we did not record ERPs, our data show an analogous behavioral signature: slower responses when expectations are violated. Hence, by leveraging the natural temporal structure of everyday events, where visual information often precedes the sounds produced by an interaction, we demonstrate that the crossmodal predictive effects are evident not only in simple artificial contexts but also in complex naturalistic events. These findings build on prior work showing that material perception is shaped by learned expectations rather than purely bottom-up cues (Alley et al., 2020; Malik et al., 2023). Whereas previous studies have demonstrated expectation-violation costs within the visual modality, the present results extend this framework to the multisensory domain.

### Material ratings reflected a weighted average of audio and video categories

When the material attribute conveyed by preceding visual information conflicted with that conveyed by incoming auditory information, participants’ ratings reflected contributions from both sources. This pattern is also consistent with the unimodal effects in rating data reported by Alley et al. (2020), where material judgments likewise shifted toward a weighted integration of prior material expectations and the available sensory evidence.

In a related crossmodal material-perception study, Fujisaki et al. (2014) examined how visual and auditory information are combined during material categorization and material-property judgments by presenting videos and impact sounds concurrently. Across a broad set of bipolar property ratings, they reported that judgments are well described by a weighted-integration account, with the relative contribution of vision versus audition varying systematically with the property being judged (e.g., greater visual influence for surface-related properties and stronger auditory influence for sound-related properties), and more comparable contributions for mechanically relevant, crossmodal attributes. The attributes we probed here align most closely with these mechanically relevant, crossmodal properties. Despite the difference in our paradigm (temporal segregation versus concurrent presentation of audiovisual information), our data also resemble a weighted-integration profile.

However, when we look more closely at the participant-level data from the current study, we see individual variation: inter-observer consistency was moderate (ICC = 0.41; Fig. 4), and between-participant random effects in prior pull accounted for a large share of the explainable variance. Thus, although the aggregate data resemble a weighted-integration profile, the relative weights varied widely across individuals: some participants relied more strongly on audition, others were more influenced by the visual prior, with many integrating both cues to varying degrees. Such pronounced inter-individual differences are not unusual in perception, particularly when the sensory input is ambiguous or permits multiple interpretations. A well known example is *The Dress*, where the same image gave rise to strikingly different percepts across observers, apparently because viewers adopted different assumptions about the scene’s illumination (Wallisch, 2017). Likewise, observers in the present task may have differed in the strength or nature of the priors they carry for the audiovisual event. One plausible source of this variability is visual sampling: eye-movement research shows that observers differ systematically, and often stably, in where they look (Bargary et al., 2017; Henderson & Luke, 2014), and such differences can have important perceptual consequences. In audiovisual speech, for example, individual differences in face viewing behavior predict the magnitude of multisensory gain, suggesting that the way visual information is sampled can shape crossmodal weighting (Rennig et al., 2020). By analogy, participants in the present task may have differed in the extent to which they sampled and encoded diagnostic pre-impact visual cues, leading some to form stronger material predictions and others to rely more heavily on the incoming sound. Differences in audiovisual binding tendencies may have contributed further, as individuals vary in temporal binding and in sensitivity to audiovisual temporal correlations, both of which could influence whether the sound is interpreted as belonging to the previewed object (Stevenson et al., 2012). Finally, although instructions were identical, task set and attention need not be uniform: participants may differ in how they construe the goal of the judgment and in which modality they attend most, even without explicit cue-directed instructions (Talsma et al., 2010). Thus, the large individual differences in prior pull may reflect meaningful variation in how observers form, maintain, and apply visually induced expectations during subsequent auditory judgments. Future work should test the basis of such differences directly.

### Effect of attribute on the magnitude of prior pull

Although most of the explainable variance in prior pull was captured by between-participant differences, a smaller portion was accounted for by the fixed effects. These fixed-effect estimates should nevertheless be interpreted with care. To keep the model parsimonious and to target cue conflict, we collapsed each bipolar attribute into two poles and restricted the analysis to contrasting incongruent pairs. As a consequence, the specific audio-video pairs differed across attributes, meaning that differences in mean prior pull between attributes could in principle reflect idiosyncrasies of the particular stimulus pairs rather than attribute-specific mechanisms. For this reason, we do not draw any conclusions from cross-attribute differences, even though prior pull was largest for the deforming/breaking-stays intact attribute. By contrast, comparisons within each attribute are more informative because the two conflict directions were defined symmetrically (visual-high/audio-low vs. visual-low/audio-high) and were evaluated within the same attribute context.

Within attributes, we observed a clear asymmetry across conflict directions. Post-hoc contrasts showed that the auditory cue exerted a stronger influence when the sound conveyed hard (vs. soft), dry (vs. wet), or metallic (vs. non-metallic). Conversely, when the sound conveyed soft, wet, or non-metallic, while the visual event implied the opposite pole, ratings showed greater prior pull toward the visual reference. For instance in the third column of Figure 7-A, when the sound is soft, the estimated maginal mean of prior pull is more negative than when the sound is hard, indicating more reliance on the visual prior when the sound conveys a soft object. We suggest that this asymmetry reflects reliability-based weighting between the visually induced expectation and the incoming auditory evidence. It has been shown across cue-combination paradigms that cue weights shift with relative reliability. For example, when the reliabilities of visual and haptic signals are manipulated, observers shift their weighting accordingly in visuo-haptic height judgments (Ernst & Banks, 2002), and in audiovisual localization, reducing visual precision decreases visual capture and increases reliance on audition (Alais & Burr, 2004). In our stimulus set, impacts associated with hard, dry, and metallic materials may yield more distinctive (i.e., higher-fidelity) acoustic cues, whereas soft, wet, and non-metallic impacts may be more acoustically underspecified for the rated attribute. This would naturally produce the observed asymmetry: stronger auditory dominance when the sound is more diagnostic, and greater prior pull when the sound provides weaker evidence.

### A Bayesian perspective on crossmodal expectation

Our findings are consistent with a Bayesian, or more broadly, a predictive-processing account of perception, in which observers combine prior expectations with incoming sensory evidence to infer the most likely cause of an event (Friston, 2005; Summerfield & de Lange, 2014). Note, however, that we do not attempt to formally model this inference here; we use the framework as an interpretive account of the observed pattern of results.

Based on our knowledge and previous experience with objects and materials, we learn to associate objects with particular material properties, and these associations often extend across modalities (Ernst & Banks, 2002; Lederman et al., 1986). For instance, a spoon-shaped object with a rigid, shiny appearance approaching the hard floor naturally sets up an expectation of a brief, high-frequency “clink” sound with a sharp onset. Therefore, in our paradigm, simply seeing an object approaching the ground elicits a prior about its material properties and, by extension, what the impact is going to sound like. The subsequent impact sound then provides the crossmodal likelihood, that is, auditory evidence about the material properties of the object. The observer’s final judgment can be described as a posterior estimate that reflects an uncertainty-weighted compromise between these two sources of information (Knill & Pouget, 2004).

This framework provides a common interpretation for the two main signatures observed in the current study. First, slower responses on incongruent trials can be interpreted as a cost of model updating: when the auditory evidence does not match the visually induced prior, large prediction errors occur, and additional processing is required to reconcile the mismatch, revise the inferred cause, or decide whether the sound should be attributed to the previewed object at all (De Loof et al., 2016; Urgen & Boyaci, 2021). Second, the same framework captures the rating data. If we treat the visual category rating implied by the preceding visual information as the prior mean and the auditory category rating implied by the sound as the likelihood mean, then the participant’s rating reflects their posterior mean, shifted toward whichever source is more precise. Crucially, this framework also offers a principled account of the large inter-individual differences in cue weighting. Weights are determined by relative precision (inverse variance). Between-participant variability in “prior pull” can therefore arise from at least three non-exclusive sources: (i) differences in the precision of the visual prior (how strongly the preview constrains expected material outcomes), (ii) differences in auditory likelihood precision (how diagnostic the impact sounds are for a given observer), and (iii) differences in causal inference, the probability that the sound and the previewed object share a common cause (Körding et al., 2007; Shams & Beierholm, 2010). Notably, the second component, differences in auditory likelihood precision, could also help explain the attribute asymmetries we observed: When the auditory cue conveys hard, dry, or metallic materials, the likelihood may be sharper (higher precision) because these impacts often contain distinctive spectral-temporal structure. In contrast, when an auditory cue conveys soft, wet, or non-metallic sounds, the acoustic evidence for the attribute being judged may be weaker, and hence the likelihood is comparatively broader. A sharper likelihood should pull the posterior strongly towards itself than the broader; which is consistent with the pattern we observed. However, future work will be needed to formalize this account by explicitly modeling priors, likelihoods, and common-cause assumptions within observers, and to test whether these parameters predict both individual weighting patterns and RT costs.

### Conclusions

In conclusion, visually derived priors about material properties of an object generate higher-level expectations about its future material state, and the influence of these expectations extends across modalities. More broadly, these top-down crossmodal effects provide a plausible mechanism for multisensory facilitation in material perception. Further understanding how such high-level crossmodal expectations are integrated with incoming sensory evidence may help explain how the perceptual system achieves efficient material perception in naturalistic settings.

## Acknowledgements

This work was supported by the Deutsche Forschungsgemeinschaft (German Research Foundation, DFG) under Germany’s Excellence Strategy (EXC 3066/1 “The Adaptive Mind”, Project No. 533717223) and by the European Union’s Horizon Europe research and innovation programme under the Marie Skłodowska-Curie grant agreement No 101226908 (EXPLORA).

## Supplementary Information

### Supplementary Figures

**Figure S1.**
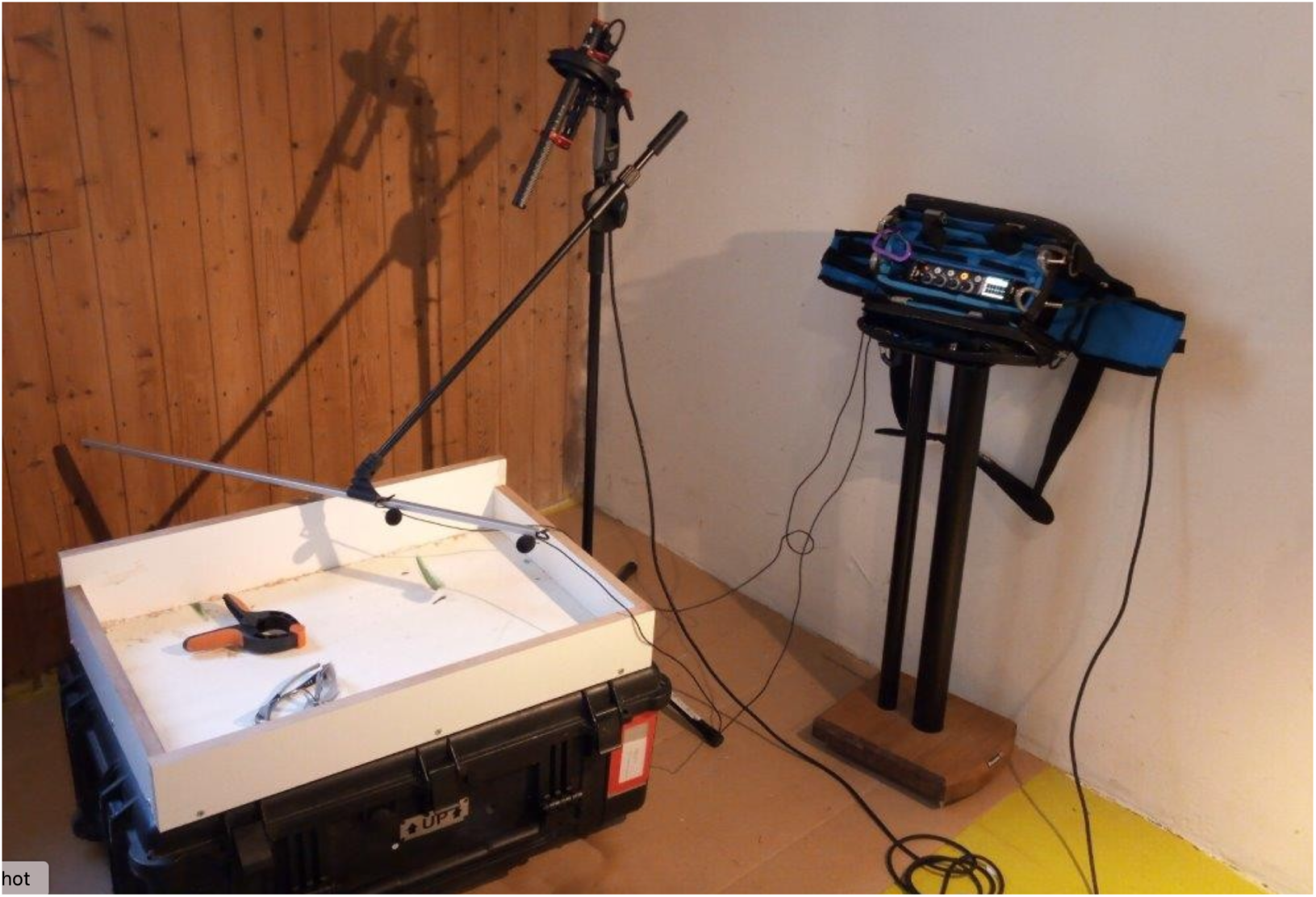
Recording setup used to capture material-specific impact sounds.

**Figure S2.**
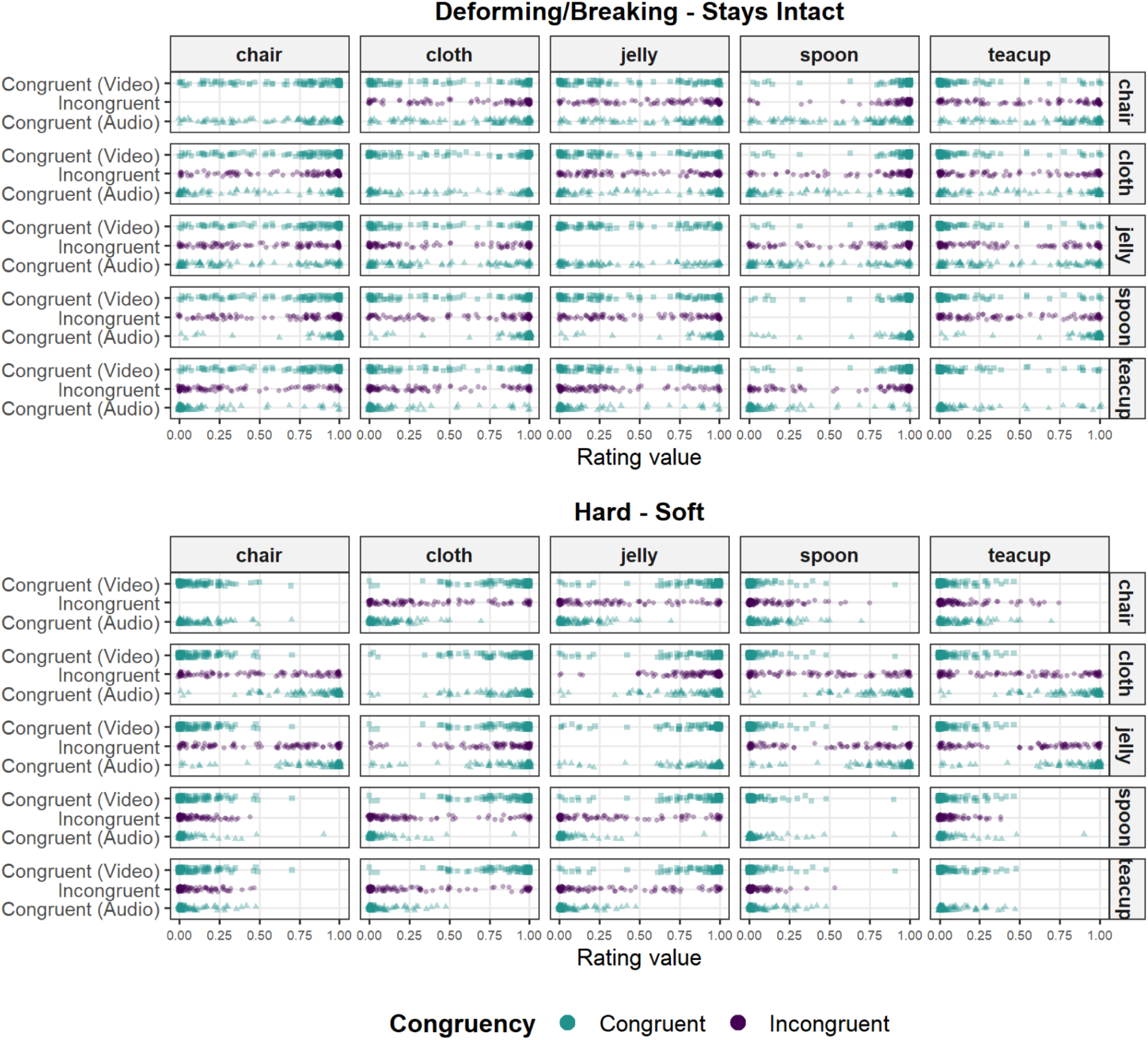
The upper and lower panels show raw ratings for the deforming/breaking-stays intact and hard-soft attributes, respectively. Each point shows an individual rating response for a given condition. Facets show the reference audio material by row and the reference video material by column. The y-axis groups ratings for each incongruent audio-video pair with the corresponding congruent audio reference and congruent video reference ratings..

**Figure S3.**
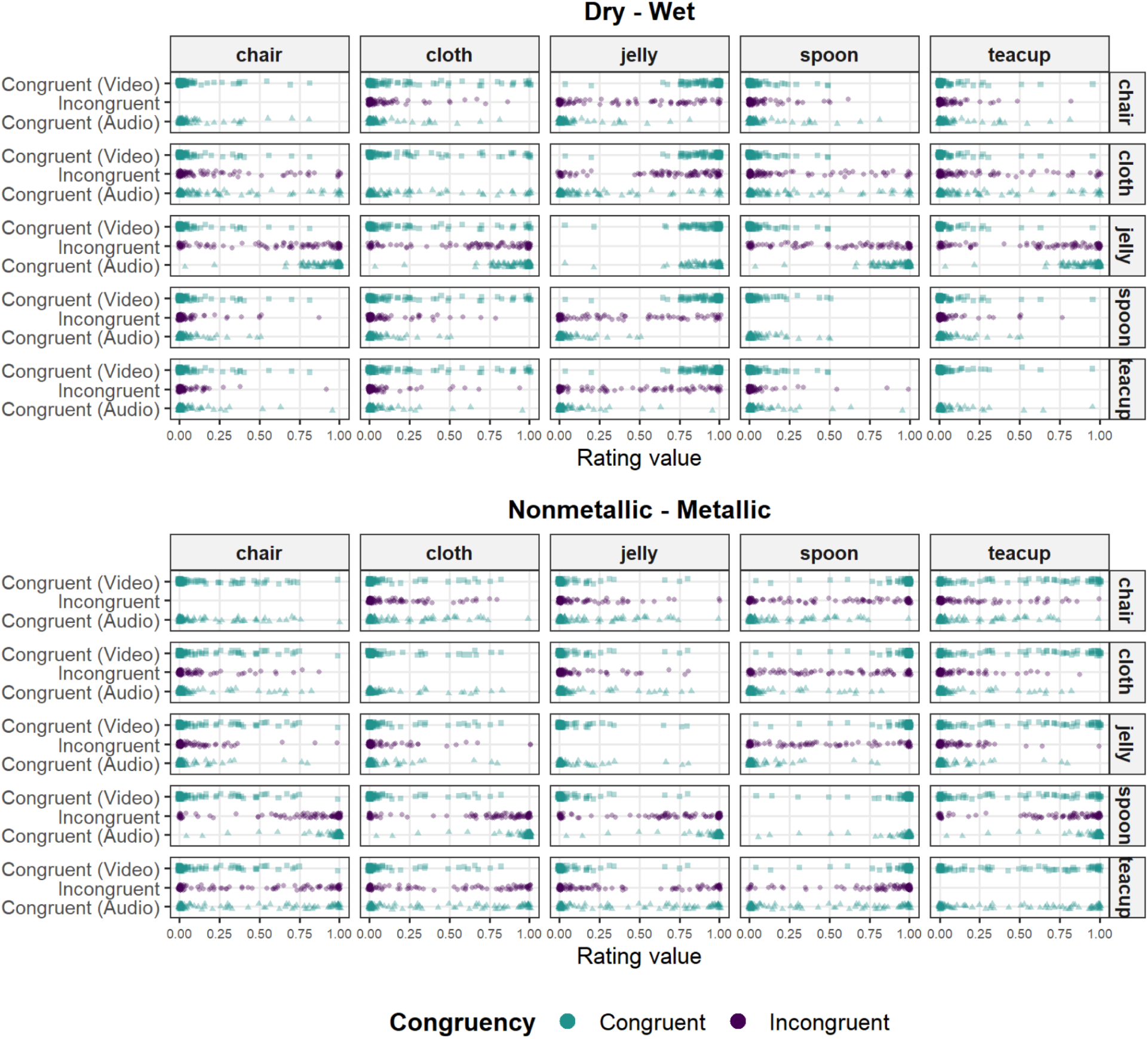
The upper and lower panels show raw ratings for the dry-wet and metallic-nonmetallic attributes, respectively. Each point shows an individual rating response for a given condition. Facets show the reference audio material by row and the reference video material by column. The y-axis groups ratings for each incongruent audio-video pair with the corresponding congruent audio reference and congruent video reference ratings.

### Supplementary Tables

**Supplementary Table S1.** Visual stimulus objects and corresponding sound-source materials.

| Material category | Visual object (stimulus) | Characteristic Sound | Sound source material/object (recording) |
| --- | --- | --- | --- |
| **shattering** | Claypot | Shatter | Terracotta clay pots, approx. 23 × 20 cm (IKEA) |
|  | Glass | Shatter | Assorted wine glasses (IKEA) |
|  | Teacup | Shatter | Thin-walled dessert bowl (IKEA) |
| **splashing** | Honey | Splash | Soy yogurt |
|  | Milk | Splash | Milk and water |
|  | Water | Splash | Water |
| **nondeforming** | Chair | Thud | Oak-wood chair |
|  | Key | Clink | Key |
|  | Spoon | Clink | Small teaspoon |
| **wobbling** | Custard | Plop | Gelatin dessert |
|  | Red Jelly | Plop | Gelatin dessert |
|  | Green Jelly | Plop | Gelatin dessert |
| **wrinkling** | linen curtain | Rustle | Linen cloth |
|  | Silk curtain | Rustle | Silk cloth |
|  | Velvet curtain | Rustle | Velvet cloth |

*IKEA and Netto refer to the retailers from which some items were sourced.*

**Supplementary Table S2.** Mapping between audio and video categories in congruent (expected) and incongruent (surprising) conditions for experiment 1, adapted from Alley et al., 2020.

| Material category | Expected/Congruent | Surprising/Incongruent | Corresponding Sound |
| --- | --- | --- | --- |
| **Shattering** | Claypot | Custard | Shatter |
|  | Glass | Linen curtain | Shatter |
|  | Teacup | Water | Shatter |
| **Splashing** | Honey | Chair | Splash |
|  | Milk | Claypot | Splash |
|  | Water | Silk curtain | Splash |
| **Nondeforming** | Chair | Milk | Thud |
|  | Key | Red jelly | Clink |
|  | Spoon | Velvet curtain | Clink |
| **Wobbling** | Custard | Glass | Plop |
|  | Red Jelly | Honey | Plop |
|  | Green Jelly | Key | Plop |
| **Wrinkling** | Linen curtain | Green jelly | Rustle |
|  | Silk curtain | Spoon | Rustle |
|  | Velvet curtain | Teacup | Rustle |

**Supplementary Table S3.**
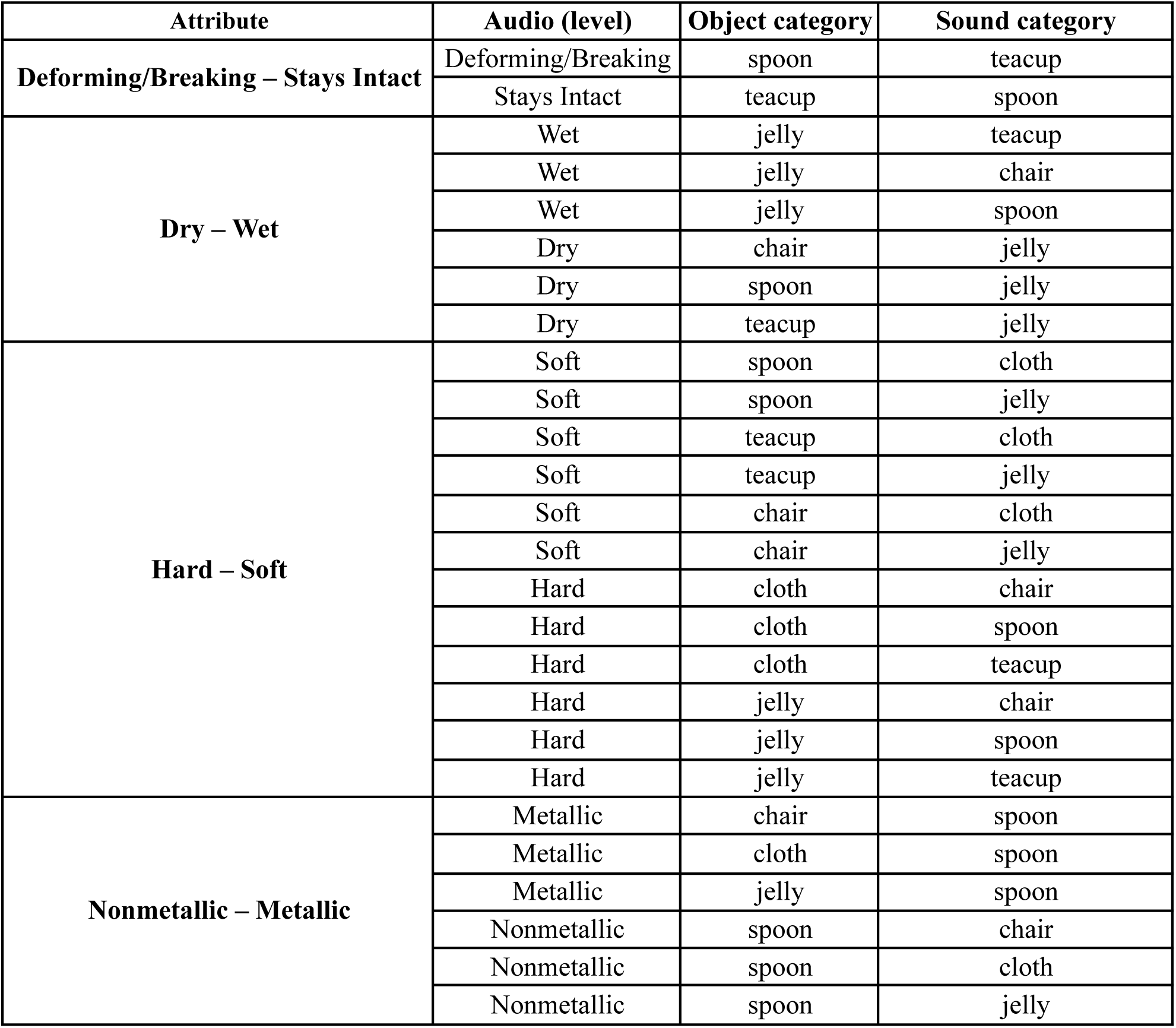
Stimulus labels for strongly contrasting incongruent audiovisual pairs, categorized by conflict direction.

